# Imagining and reading actions: towards similar motor representations

**DOI:** 10.1101/2022.06.07.495123

**Authors:** W Dupont, C Papaxanthis, C Madden-Lombardi, F Lebon

## Abstract

While action language and motor imagery both engage the motor system, determining whether these two processes indeed share the same motor representations would contribute to better understanding their underlying mechanisms. We conducted two experiments probing the mutual influence of these two processes. In Exp.1, hand-action verbs were presented subliminally, and participants (n=36) selected the verb they thought they perceived from two alternatives. When congruent actions were imagined prior to this task, accuracy significantly increased, i.e. participants were better able to “see” the subliminal verbs. In Exp.2, participants (n=19) imagined hand flexion or extension, while corticospinal excitability was measured via transcranial magnetic stimulation. Corticospinal excitability was modulated by action verbs subliminally presented prior to imagery. Specifically, the typical increase observed during imagery was suppressed after presentation of incongruent action verbs. This mutual influence of action language and motor imagery, both at behavioral and neurophysiological levels, suggests overlapping motor representations.

## Introduction

Comprehending described events is one of the most sophisticated functions of human cognition. Over the past several decades, a growing body of literature has provided support for an embodied view of language comprehension, suggesting that understanding language evokes simulations in the systems of the brain that are used for perception and action^1–3^. According to proponents of this idea, action language would automatically and unconsciously solicit a motor representation in order to visualize features of described action, such as direction and duration ^4–6^, as well as the effector used ^7,8^. Some researchers postulate that these motor representations serve to more efficiently understand the action described ^4,9,10^. Consistent with these ideas, the comprehension of described actions is often accompanied by changes in motor output ^11–17^, characterized by the increase of corticospinal excitability in Transcranial Magnetic Stimulation (TMS) studies ^18–21^ or the involvement of motor areas in neuro-imaging studies ^22–26^.

This automatic, implicit, and unconscious motor representation approaches a well-known process called motor imagery, which is the explicit mental simulation of action without concomitant movement ^27^. During motor imagery, many TMS and imaging studies reveal an increase of corticospinal excitability ^28–32^ and activation of the motor network overlapping with that observed during actual movements ^33–35^. These neurophysiological substrates would reflect the elaboration of motor representations to form mental motor images ^27,36^.

Nevertheless, few studies have directly compared the motor representations engaged during motor imagery and action reading. According to some authors, motor representations triggered during action verb reading correspond to motor imagery in an unconscious form ^37,38^. This raises questions about the neurophysiological similarity of these two processes. Yang et Shu (2014)^39^ report an overlap of brain areas during motor imagery and passive action verb reading. However, a contradictory finding is reported by Willems et al. (2010)^40^, who describe these two processes as distinct and engaging different motor representations. Given these results, it remains unclear whether action language and motor imagery consist of separate processes, or whether they rely on the same cognitive process with varying levels of motor network involvement.

The present paper aims to shed new light on this issue by exploring the mutual influence of action reading and motor imagery. To assess automatic language representations (without any imagery strategies), we opted to use subliminal priming, which is a well-known paradigm in cognitive neuroscience ^41–43^. In a typical subliminal priming experiment, words are preceded and followed by masking patterns, making them invisible. Behavioral priming experiments have repeatedly shown that subliminal words are nevertheless processed unconsciously ^44–48^ and have even shown interference with the simultaneous preparation and subsequent execution of an arm extension movement ^49^. In a first behavioral experiment, we probed whether motor imagery could facilitate access to motor representations during action reading, consequently helping the perception of subliminal action verbs. In a second neurophysiological experiment, we used TMS to test whether the presentation of congruent and incongruent subliminal action verbs would modulate the corticospinal excitability increase classically observed during motor imagery. If the representations generated during action word reading and motor imagery do overlap (at least partially), we would expect to see this bidirectional influence.

## Results and Discussion

### Behavioral Experiment: influence of motor imagery on subliminal action reading

In experiment 1, thirty-six participants imagined a manual flexion or extension action before the subliminal presentation of a congruent or incongruent action verb (or words unrelated to action). Then, the participants were instructed to select the verb they thought they perceived from two alternatives, as in a forced choice task (e.g.,”I squeeze” or “I extend”, see Fig.1). We measured the percentage of correct responses and reaction times.

**Figure 1:**
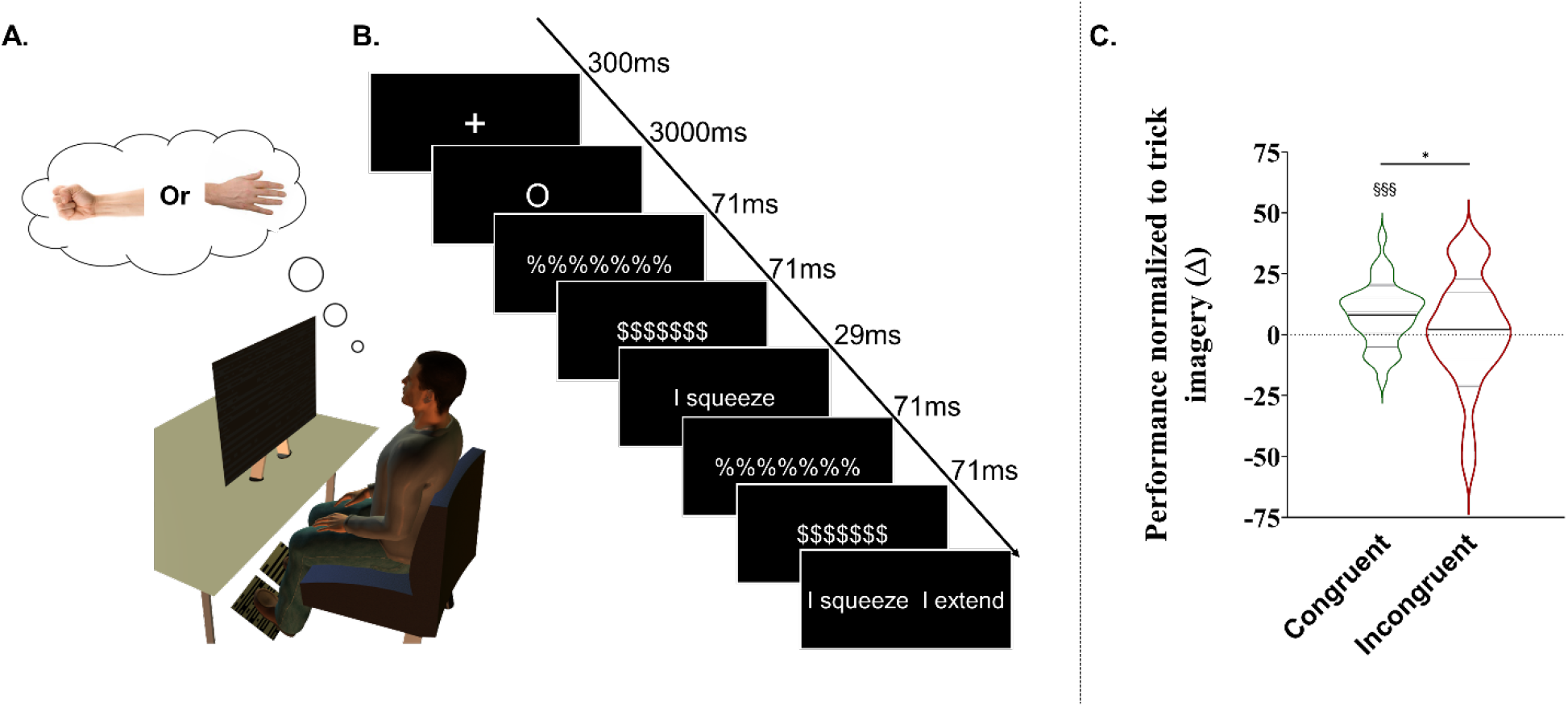
**A**. Illustration of the procedure, with the position of the participant during the subliminal reading and motor imagery tasks. The participants imagined a hand flexion and extension or they did not imagine any movements (Control trials). **B**. Illustration of the display for one trial. A trial started with the attentional cross (duration= 300 ms), followed by the motor imagery signal (3000 ms), two successively masks (71 ms each; Mask 1: %%%%%% and Mask 2: $$$$$$), the subliminal stimuli (29 ms), the two masks, and the two alternatives (forced choice task). On congruent trials the subliminal action word is congruent with the previously imagined action (imagined flexion – “squeeze” presented), whereas on Incongruent trials the subliminal action word is incongruent with the previously imagined action (imagined extension – “squeeze” presented). On trick trials a word is subliminally presented that is unrelated to the previously imagined flexion or extension action (imagined flexion – “people” presented), but participants still had to select from the two (unpresented) verbs as in experimental trials (“squeeze” or “extend”). **C**. Violin plots represent normalized performance (Δ on Trick trials). Thick and thin horizontal lines mark mean and SD, respectively. ANOVA revealed a congruence effect *=p<0.05. §§§=p<0.001 indicates a significant difference from zero (Trick trials).

In Congruent trials, the subliminal action word was congruent with the previously imagined action (imagined flexion – “squeeze” presented), whereas in Incongruent trials, the subliminal action word was incongruent with the previously imagined action (imagined extension – “squeeze” presented). In Trick trials, a word that was unrelated to the previously imagined flexion or extension action was subliminally presented, but participants still had to select from the two (unpresented) verbs as in experimental trials. These trials were included to assess whether participants used a strategy in which they chose the verb that best matched the imagined action. Finally, we used Control trials (without imagery) to ensure that the word presentation was indeed subliminal. For these trials, the percentage of correct responses (52.69 ±11.06%) did not differ from chance, i.e. 50% (p=0.154).

If motor imagery can influence subliminal action reading, we expect to observe an increase in correct responses in Congruent trials (or a decrease in correct responses for Incongruent trials) in comparison to Trick trials. We analyzed the percentage of correct responses in Congruent and Incongruent trials normalized to that of Trick trials (Δ). Results of reaction times are presented in Supplementary section.

We performed a repeated measures ANOVA with Congruence (congruent/incongruent) and Verb type (extension/flexion) as within-subject factors. We observed a main effect of Congruence (F_1,35_=5.150, p=0.029, ηp^2^=0.128), with better performance for Congruent (8.26 ±12.51%) than Incongruent trials (1.78 ±22.46%). We did not observe a significant effect of Verb type (F_1,35_=0.998, p=0.324, ηp^2^=0.027) nor an interaction between Congruence and Verb type (F_1,35_=2.722, p=0.107, ηp^2^=0.072) (See Figure 1). These results demonstrate that motor imagery can help in perceiving subliminal action verbs, most likely by facilitating access to motor representations during subconscious action reading.

Also, one-sample t-tests showed that Congruent (p <0.001) but not Incongruent (p=0.636) trials differed from zero, i.e., Trick trials. During Trick trials, the participants imagined an action but non-related action verbs were subliminally presented. This result indicates that the congruent association between motor imagery and action verb renders the subliminal protocol less subliminal. The greater percentage of correct responses during Congruent trials (60.16 ±10.52%) shows that the participants did not just pick the action verb they just imagined, but they “perceived” the subliminal action verb that was presented (See Supplementary section for % of correct responses in all conditions).

The fact that motor imagery is able to influence lexical access of these action words suggests that these two processes indeed share motor representations.

### Neurophysiological Experiment: influence of subliminal action reading on motor imagery

In experiment 2, we flipped the order of presentation within the trials, such that the action verb was presented subliminally before participants (n=19) imagined the manual flexion or extension that was either congruent or incongruent with the previously presented verb.

Single-pulse TMS were delivered over the finger/hand muscle area of the left primary motor cortex during motor imagery. Corticospinal excitability was assessed in the form of motor-evoked potentials (MEPs) amplitude. In order to probe for muscle-specific effects, we recorded MEPs in two muscles involved in flexion movements (Flexor Digitorum Superficialis, Flexor Carpi Radialis) and two muscles involved in extension movements (Extensor Digitorum Superficialis, Extensor Carpi Radialis). We analyzed MEPs for which the muscle action matched the imagined action, i.e., MEPs of FDS and FCR when imagining flexion and MEPs of EDS and ECR when imagining extension.

In Control Imagery trials, participants imagined flexion or extension actions after a chain of meaningless consonants was subliminally presented. We added Control trials without imagery (and without TMS) at the end of the experiment to ensure that the verb presentation was indeed subliminal (50.69 ±10.15%, p=0.767). Congruent, Incongruent and Control Imagery trials were normalized to MEPs recorded at rest, i.e., without imagery nor subliminal verbs. If subliminal action reading can influence motor imagery, we expect to observe a modulation of corticospinal excitability in Congruent and Incongruent trials in comparison to the Control Imagery trials, for all tested muscles.

The overall ANOVA revealed a main effect of Congruence (F_2,36_=5.441, p=0.008, ηp^2^=0.232), which was similar for all muscles, as we did not find any main effect of Muscle (F_3,54_=1.086, p=0.362, ηp^2^=0.056) nor a Congruence by Muscle interaction (F_6,108_=0.789 p=0.579, ηp^2^=0.042) (See Figure 2). Paired comparisons with Tukey corrections showed larger MEP amplitudes for Congruent (33.07 ±37.37%, p=0.012, Cohen’s d= 2.80) and Control Imagery trials (30.35 ±47.96%, p=0.031, Cohen’s d= 2.09) in comparison to Incongruent trials (11.49 ±31.23%). We did not observe any difference between Congruent and Control Imagery trials (p=0.923, Cohen’s d= 0.28).

**Figure 2:**
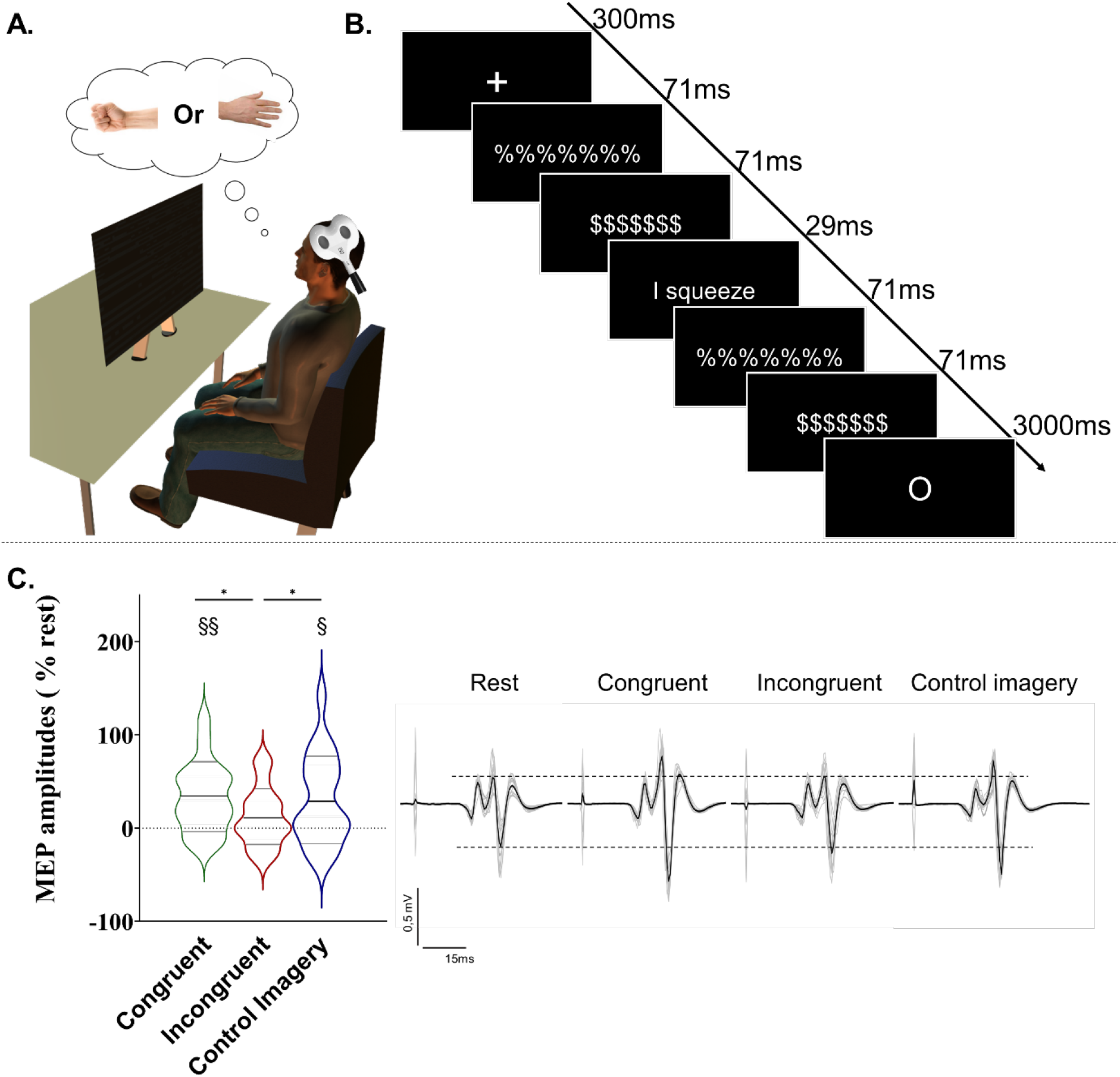
**A**. Illustration of the procedure, with the position of the participant during the subliminal reading and motor imagery tasks. Participants imagined either a hand flexion, an extension or they did not imagine any movements (Control without imagery and rest trials). The TMS coil was positioned over the primary motor cortex **B**. Illustration of the display for one trial. A trial started with the attentional cross (duration= 300 ms), followed by two successively masks (71 ms each; Mask 1: %%%%%% and Mask 2: $$$$$$), the subliminal stimuli (29 ms), the two masks, and the motor imagery signal (3000 ms). TMS pulses were delivered at 300, 350 or 400ms after the motor imagery signal. Congruent and Incongruent trials correspond to flexion or extension imagination preceded by a subliminal presentation of a congruent or incongruent action verb, respectively. Control Imagery trials correspond to flexion or extension imaginations after a subliminal presentation of a chain of meaningless consonants (e.g.,”tjgkdl”). **C**. Violin plots represent normalized MEPs (% rest). Thick and thin horizontal lines mark mean and SD, respectively. ANOVA revealed a congruence effect *=p<0.05. The § symbol indicates a difference from rest. The right side of the panel illustrates raw MEPs of a typical subject (grey lines). The black line is the average MEP of the condition for this participant.

One-sample t-tests on normalized MEPs (% rest) yielded significant differences from rest for Congruent (p=0.001) and Control Imagery (p=0.012) but not for the Incongruent trials (p=0.126). These results demonstrate that language about actions can influence our ability to imagine hand movements. It is noteworthy that the visual presentation of action verbs, albeit subliminal, was indeed able to modulate the motor system, yielding a measurable difference at hand muscles. We confirmed the classic increase of corticospinal excitability during motor imagery ^28–32,50^ (Control Imagery trials). Although the subliminal presentation of action verbs congruent with this imagination produced no additional increase, this increase was suppressed when incongruent action verbs were subliminally presented before imagination.

Taken together, the results of these two experiments highlight a mutual influence of motor imagery and action reading at both behavioral and neurophysiological levels. In Exp.1, percentages of correct responses suggested that participants were better able to “see” the subliminally presented word if it was congruent rather than incongruent with the action that had been imagined at the start of the trial. We interpret these results as facilitation for lexical access to the action verb when the corresponding motor representation is already activated or primed from the preceding motor imagery, although it is also possible that inhibition occurs when a competing motor representation is already activated or primed from the preceding incongruent motor imagery. These original results feed the link between cognition and action, a relevant step in the comprehension of previous studies suggesting shared brain activation patterns between motor imagery and action language ^39^ or perception ^51–54^. This reminds interactions at the perceptual level where conscious perception may be influenced by mental visual imagery ^55–64^.

In Exp.2, we flipped the order of presentation within the trials, such that the action verb phrase was presented subliminally before the participant imagined the manual flexion or extension. The subliminal presentation appeared as a visual blip before the cue to imagine the action, and participants were usually not aware that a verb had even been presented. However, congruent action verbs increased excitability at hand muscles compared to rest, while incongruent action verbs suppressed this increase, yielding a significant difference between these two conditions. This second experiment provides first evidence that subliminal reading of action verbs modulates the neurophysiological markers of motor imagery. To note that congruent action verbs did not increase to a greater extent the corticospinal excitability while imagining without action verbs ^28–32,50^. This may be explained by i) a ceiling effect, i.e., MEPs amplitude cannot further increase during motor imagery, ii) an absence of congruent priming effect on corticospinal excitability or iii) an inhibitory mechanism observed during motor imagery ^65–67^, which prevents any extra increase with subliminal priming.

The results from these two experiments support the idea that motor imagery and action language can influence each other at both the neurophysiological and behavioral levels, and thus it seems quite likely that these two processes share motor representations.

## Material and method

### Experiment 1

#### Participants

Forty-one healthy right-handed individuals (19 females; mean age = 23.92 years-old; range 18-35) participated in the experiment. Participants’ handedness was assessed by the Edinburgh inventory ^68^. All subjects were French native speakers without neurological, physical and psychiatric pathology.

#### Stimuli

Ten hand action verbs were selected for the experiment, half describing finger and wrist extension (e.g., “I extend”), and half describing flexion of these joints (e.g., “I squeeze”). These verbs always appeared in the first person present tense. Using the Lexique.org database ^69^, we controlled various psycholinguistic factors between the two sets of verbs (written frequency, number of characters, number of syllables and spelling neighbors; see Table 1 for details).

The Trick trials (Control Imagery condition) consisted of five words that did not evoke movement (e.g., “people”; see Supplementary section for details). This allowed us to identify participants who employed a strategy in which they systematically selected the verb they just imagined.

#### Procedure

Participants sat in an armchair while stimuli were presented on a 19-inch LCD monitor by a home-made software, which also recorded behavioral responses and electromyographic activity (10-1000 Hz, Biopac Systems Inc.). Throughout the recording, participants were instructed not to move their hands while they imagined movements followed by subliminal stimuli. We adapted the subliminal paradigm of Dehaene et al.^42^ with two successively displayed pattern masks (duration=71ms; Mask 1: %%%%%% and Mask 2: $$$$$$), the subliminal stimuli (duration=29ms), and again the same masks. At the end of the trial, two words were presented and the participant had to choose which one he/she thought he/she had perceived in the preceding subliminal presentation (forced choice subliminal task). In order to avoid interference with motor imagery of hand actions, the choice was made by pressing pedals with the feet. Each trial started with a fixation cross, followed by a signal indicating to imagine hand action, then by the subliminal paradigm before the forced choice subliminal task (See Figure 1).

A familiarization session was conducted before the experimental session, in which participants saw eight trials. The experimental session was divided into six blocks with motor imagery and one without (Control task). Each motor imagery block included 30 imagined trials, yielding 180 imagined trials total in the experiment. For half of the blocks, the subject imagined a wrist and finger flexion movement. Conversely, for the remaining half, the subject imagined a wrist and finger extension. For example, in the three flexion imagery blocks, extension action verbs (e.g., “I extend”), flexion action verbs (e.g., “I squeeze”) and control words (e.g., “screen” or “people”) were presented subliminally after the motor imagery, corresponding respectively to the Incongruent, Congruent and Trick trials. There were the same conditions for the extension imagery. Block order was randomized and counterbalanced.

#### Data and statistical analysis

We measured the percentage of correct responses at the forced choice subliminal task, i.e. when the participants selected the verb that was indeed subliminally presented. In Trick trials, although neither of the two choices had been presented, we considered a response “correct” when the action verb just imagined was selected (only non-related action verbs were presented). To eliminate imagery strategies, we excluded participants that nearly always (≥85%) selected the verb that matched the imagined action in Trick trials (not reflecting a possible effect of real verb perception). This resulted in the exclusion of 5 participants.

Then, performance (% correct responses) for Congruent and Incongruent trials was normalized to Trick trials (Δ). Reaction times (RT) for the forced choice task reflected the time between the presentation of the two alternatives on the screen and the pedal press. RTs in experimental conditions were normalized to the mean of RTs on Trick trials (see Supplementary section). Statistics and data analyses were performed using the Statistica software (Stat Soft, France). Normality and sphericity of the data were checked with Shapiro-Wilk and Mauchly tests, respectively. The data are presented as mean values (±standard deviation) and the alpha value was set at 0.05.

### Experiment 2

#### Participants

Twenty healthy right-handed individuals (9 women; mean age = 22.57 years-old; range 18-28) participated in the experiment. Participants’ handedness was assessed by the Edinburgh inventory ^68^. All subjects were French native speakers without neurological, physical and psychiatric pathology. Volunteers confirmed their participation with written consent at a medical visit before the TMS protocol. The local Ethics Committee approved experimental protocol and procedures in accordance with the Declaration of Helsinki (CPP 2017-A00064-49).

#### Stimuli and procedure

Stimuli were identical to Experiment 1, except that as a Control Imagery condition we used five chains of meaningless consonant letters (unpronounceable in French; e.g., “tjgkdl”; see Supplementary section for details), allowing comparisons of Congruent and Incongruent trials to this Control Imagery trials without action verbs.

The procedure was almost identical to Experiment 1, except for two features. First, participants performed the motor imagery at the end of the trial, just after the subliminal paradigm. Second, we used TMS to probe corticospinal excitability at rest, as well as various stimulation delays during the motor imagery task (300, 350, and 400ms after the onset of the signal to imagine). These latencies were chosen in order to allow enough time for the subject to imagine while remaining in the optimal window of investigating priming effect (0-500ms).

A familiarization session was conducted before the experimental session, in which participants performed ten trials, each starting with a fixation cross, followed immediately by two successively displayed pattern masks, the subliminal stimuli, and again the same masks before the motor imagery signal.

The experimental session was divided into six blocks with motor imagery and one without (Rest condition). Each motor imagery block included 15 imagined trials, yielding 90 imagined trials total in the experiment. For half of the blocks, the subject imagined a wrist and finger flexion movement. Conversely, for other blocks, the subject imagined a wrist and finger extension. For example, in the three imagined extension blocks, Congruent Action verbs (e.g., “I extend”), Incongruent Action verbs (e.g., “I squeeze”) and Control Imagery (e.g., “tjgkdl”) were presented subliminally in three separate blocks. Block order was randomized and counterbalanced.

#### Transcranial magnetic stimulation

Single-pulse TMS was generated from an electromagnetic stimulator Magstim 200 (Magstim Company Ltd, Whitland) and using a figure-eight coil (70 mm in diameter). The coil was placed over the contralateral left hemisphere to target the motor area of the Extensor and Flexor Digitorum Superficialis muscles (EDS and FDS respectively) and the Extensor and Flexor Radialis Carpi (ECR and FCR respectively) muscles of the right forearm. First, we individually determined the precise stimulation site (hotspot), where the MEP amplitude at the four muscles was the highest and the most consistent. Then, the resting motor threshold of each participant was determined as the minimal TMS intensity necessary to induce a MEP of 0.05mV peak-to-peak amplitude for 5 trials out of 10. During the experimental session, TMS pulses were delivered at 120% of the resting motor threshold.

#### EMG recording

The EMG signal was recorded by 10mm-diameter surface electrodes (Contrôle Graphique Médical, Brice Comte-Robert, France) placed over the FDS, EDS, FCR and ECR muscles of the right forearm. In order to reduce the noise in the EMG signal (< 20μV), the skin was shaved and cleaned. The EMG signals were amplified and bandpass filtered on-line (10-1000 Hz, Biopac Systems Inc.) and digitized at 2000 Hz for off-line analysis. We measured the EMGrms signal from each muscle.

#### Data and statistical analysis

EMG data were extracted with Matlab (The MathWorks, Natick, Massachusetts, USA) and we measured peak-to-peak MEP amplitude. Data falling 2.5 SDs above or below individual means for each experimental condition were removed before analysis (1.63%). Then, the average MEP amplitude for each condition (Congruent, Incongruent and Control Imagery) was normalized in comparison to Rest condition (%). One participant was removed from the final analysis due to extreme values. To ensure that MEP amplitudes were not contaminated by muscular pre-activity, we compared with an ANOVA the EMGrms (100 ms window before the TMS artifact) between the experimental and Rest conditions (see Supplementary section). Statistics and data analyses were performed using the Statistica software (Stat Soft, France). Normality and sphericity of the data were checked with Shapiro-Wilk and Mauchly test, respectively. The data are presented as mean values (±standard deviation) and the alpha value was set at 0.05.

## Supporting information

e.g. Supplemental section

## Author Contributions

Experiment design: WD, CML, FL

Data collection: WD

Statistical analysis: WD, CML, FL

Manuscript preparation: WD, CP, CML, FL

## Data availability statement

All data from this study are available at https://osf.io/fzmt6/?view_only=3869185be8c34183bf3d6b94727a0dbb

## References

1. Barsalou, L. W. Grounded Cognition. Annu. Rev. Psychol. 59, 617–645 (2008).

2. Fischer, M. H. & Zwaan, R. A. Embodied language: A review of the role of the motor system in language comprehension. Q. J. Exp. Psychol. 61, 825–850 (2008).

3. Gallese, V. & Lakoff, G. The brain’s concepts: The role of the sensory-motor system in conceptual knowledge. Cognitive Neuropsychology vol. 22 455–479 (2005).

4. Glenberg, A. M. & Kaschak, M. P. Grounding language in action. Psychon. Bull. Rev. 9, 558–565 (2002).

5. Kaschak, M. P. et al. Perception of motion affects language processing. Cognition 94, (2005).

6. Matlock, T. Fictive motion as cognitive simulation. Mem. Cogn. 32, 1389–1400 (2004).

7. Bergen, B., Narayan, S. & Feldman, J. UC Merced Proceedings of the Annual Meeting of the Cognitive Science Society Title Embodied Verbal Semantics: Evidence from an Image-Verb Matching Task Publication Date Embodied Verbal Semantics: Evidence from an Image-Verb Matching Task. escholarship.org 25 (2003).

8. Bergen, B., Chang, N. & Narayan, S. UC Merced Proceedings of the Annual Meeting of the Cognitive Science Society Title Simulated Action in an Embodied Construction Grammar Publication Date. escholarship.org 26 (2004).

9. Bailey, D. R. When Push Comes to Shove: A Computational Model of the Role of Motor Control in the Acquisition of Action Verbs. Diss. Abstr. Int. B Sci. Eng. 59, (1998).

10. Barsalou, L. W. Perceptual symbol systems. Behavioral and Brain Sciences vol. 22 577–609 (1999).

11. Andres, M., Finocchiaro, C., Buiatti, M. & Piazza, M. Contribution of motor representations to action verb processing. Cognition 134, 174–184 (2015).

12. Pulvermüller, F., Härle, M. & Hummel, F. Walking or talking?: Behavioral and neurophysiological correlates of action verb processing. Brain Lang. 78, 143–168 (2001).

13. Rabahi, T., Fargier, P., Rifai Sarraj, A., Clouzeau, C. & Massarelli, R. Effect of Action Verbs on the Performance of a Complex Movement. PLoS One 8, e68687 (2013).

14. Rabahi, T., Sarraj, A. R., Fargier, P., Clouzeau, C. & Massarelli, R. Action verb and motor performance. Kinesitherapie 12, 42–46 (2012).

15. Klepp, A., van Dijk, H., Niccolai, V., Schnitzler, A. & Biermann-Ruben, K. Action verb processing specifically modulates motor behaviour and sensorimotor neuronal oscillations. Sci. Rep. 9, 15985 (2019).

16. Taylor, L. J. & Zwaan, R. A. Motor resonance and linguistic focus. Q. J. Exp. Psychol. 61, 896–904 (2008).

17. Zwaan, R. A. & Taylor, L. J. Seeing, acting, understanding: Motor resonance in language comprehension. J. Exp. Psychol. Gen. 135, 1–11 (2006).

18. Innocenti, A., De Stefani, E., Sestito, M. & Gentilucci, M. Understanding of action-related and abstract verbs in comparison: a behavioral and TMS study. Cogn. Process. 15, 85–92 (2014).

19. Labruna, L., Fernández-Del-Olmo, M., Landau, A., Duqué, J. & Ivry, R. B. Modulation of the motor system during visual and auditory language processing. Exp. Brain Res. 211, 243–250 (2011).

20. Papeo, L., Vallesi, A., Isaja, A. & Rumiati, R. I. Effects of TMS on different stages of motor and non-motor verb processing in the primary motor cortex. PLoS One 4, e4508 (2009).

21. Papeo, L., Pascual-Leone, A. & Caramazza, A. Disrupting the brain to validate hypotheses on the neurobiology of language. Frontiers in Human Neuroscience vol. 7 (2013).

22. Hauk, O., Johnsrude, I. & Pulvermüller, F. Somatotopic Representation of Action Words in Human Motor and Premotor Cortex. Neuron 41, 301–307 (2004).

23. Van Dam, W. O., Rueschemeyer, S. A. & Bekkering, H. How specifically are action verbs represented in the neural motor system: An fMRI study. Neuroimage 53, 1318–1325 (2010).

24. Aziz-Zadeh, L., Wilson, S. M., Rizzolatti, G. & Iacoboni, M. Congruent Embodied Representations for Visually Presented Actions and Linguistic Phrases Describing Actions. Curr. Biol. 16, 1818–1823 (2006).

25. Tettamanti, M. et al. Listening to action-related sentences activates frontoparietal motor circuits. J. Cogn. Neurosci. 17, 273–281 (2005).

26. Wu, H. et al. Dissociable somatotopic representations of Chinese action verbs in the motor and premotor cortex. Sci. Rep. 3, (2013).

27. Decety, J. The neurophysiological basis of motor imagery. Behavioural Brain Research vol. 77 45–52 (1996).

28. Grosprêtre, S., Ruffino, C. & Lebon, F. Motor imagery and cortico-spinal excitability: A review. European Journal of Sport Science vol. 16 317–324 (2016).

29. Facchini, S., Muellbacher, W., Battaglia, F., Boroojerdi, B. & Hallett, M. Focal enhancement of motor cortex excitability during motor imagery: A transcranial magnetic stimulation study. Acta Neurol. Scand. 105, 146–151 (2002).

30. Tremblay, F., Tremblay, L. E. & Colcer, D. E. Modulation of corticospinal excitability during imagined knee movements. J. Rehabil. Med. 33, 230–234 (2001).

31. Fadiga, L. et al. Corticospinal excitability is specifically modulated by motor imagery: A magnetic stimulation study. Neuropsychologia 37, 147–158 (1998).

32. Rossini, P. M., Rossi, S., Pasqualetti, P. & Tecchio, F. Corticospinal excitability modulation to hand muscles during movement imagery. Cereb. Cortex 9, 161–167 (1999).

33. Hétu, S. et al. The neural network of motor imagery: An ALE meta-analysis. Neuroscience and Biobehavioral Reviews vol. 37 930–949 (2013).

34. Hardwick, R. M., Caspers, S., Eickhoff, S. B. & Swinnen, S. P. Neural correlates of action: Comparing meta-analyses of imagery, observation, and execution. Neuroscience and Biobehavioral Reviews vol. 94 31–44 (2018).

35. Lotze, M. et al. Activation of cortical and cerebellar motor areas during executed and imagined hand movements: An fMRI study. J. Cogn. Neurosci. 11, 491–501 (1999).

36. Jeannerod, M. Motor Cognition: What Actions Tell the Self. Motor Cognition: What Actions Tell the Self (2008). doi:10.1093/acprof:oso/9780198569657.001.0001.

37. Tomasino, B. & Rumiati, R. I. At the Mercy of Strategies: The Role of Motor Representations in Language Understanding. Front. Psychol. 4, (2013).

38. Tomasino, B., Fink, G. R., Sparing, R., Dafotakis, M. & Weiss, P. H. Action verbs and the primary motor cortex: a comparative TMS study of silent reading, frequency judgments, and motor imagery. Neuropsychologia 46, 1915–26 (2008).

39. Yang, J. & Shu, H. Passive reading and motor imagery about hand actions and tool-use actions: An fMRI study. Exp. Brain Res. 232, 453–467 (2014).

40. Willems, R. M., Toni, I., Hagoort, P. & Casasanto, D. Neural dissociations between action verb understanding and motor imagery. J. Cogn. Neurosci. 22, 2387–2400 (2010).

41. Dehaene, S. The neural bases of subliminal priming. Funct. neuroimaging Vis. Cogn. (attention Perform. Ser. 20 205–224 (2004).

42. Dehaene, S. et al. Cerebral mechanisms of word masking and unconscious repetition priming. Nat. Neurosci. 4, 752–758 (2001).

43. Naccache, L., Blandin, E. & Dehaene, S. Unconscious masked priming depends on temporal attention. Psychol. Sci. 13, 416–424 (2002).

44. Bowers, J. S., Vigliocco, G. & Haan, R. Orthographic, Phonological, and Articulatory Contributions to Masked Letter and Word Priming. J. Exp. Psychol. Hum. Percept. Perform. 24, 1705–1719 (1998).

45. Cheesman, J. & Merikle, P. M. Priming with and without awareness. Percept. Psychophys. 36, 387–395 (1984).

46. Ferrand, L. & Grainger, J. Effects of Orthography Are Independent of Phonology in Masked Form Priming. Q. J. Exp. Psychol. Sect. A 47, 365–382 (1994).

47. Forster, K. I. & Davis, C. Repetition priming and frequency attenuation in lexical access. J. Exp. Psychol. Learn. Mem. Cogn. 10, 680–698 (1984).

48. Marcel, A. J. Conscious and unconscious perception: Experiments on visual masking and word recognition. Cogn. Psychol. 15, 197–237 (1983).

49. Boulenger, V. et al. Subliminal display of action words interferes with motor planning: A combined EEG and kinematic study. J. Physiol. Paris 102, 130–136 (2008).

50. Lebon, F., Byblow, W. D., Collet, C., Guillot, A. & Stinear, C. M. The modulation of motor cortex excitability during motor imagery depends on imagery quality. Eur. J. Neurosci. 35, 323–331 (2012).

51. Cichy, R. M., Heinzle, J. & Haynes, J. D. Imagery and perception share cortical representations of content and location. Cereb. Cortex 22, 372–380 (2012).

52. McNorgan, C. A meta-analytic review of multisensory imagery identifies the neural correlates of modality-specific and modality-general imagery. Front. Hum. Neurosci. 6, (2012).

53. Pearson, J., Naselaris, T., Holmes, E. A. & Kosslyn, S. M. Mental Imagery: Functional Mechanisms and Clinical Applications. Trends in Cognitive Sciences vol. 19 590–602 (2015).

54. Dijkstra, N., Bosch, S. E. & van Gerven, M. A. J. Shared Neural Mechanisms of Visual Perception and Imagery. Trends in Cognitive Sciences vol. 23 423–434 (2019).

55. Pearson, J., Clifford, C. W. G. & Tong, F. The Functional Impact of Mental Imagery on Conscious Perception. Curr. Biol. 18, 982–986 (2008).

56. Chang, S., Lewis, D. E. & Pearson, J. The functional effects of color perception and color imagery. J. Vis. 13, (2013).

57. Aleman, A., D’Alfonso, A. A. L., Postma, A. & De Haan, E. H. F. Involvement of primary visual cortex in visual mental imagery: A transcranial magnetic stimulation study. Neurosci. Res. Commun. 25, 25–31 (1999).

58. Broggin, E., Savazzi, S. & Marzi, C. Similar effects of visual perception and imagery on simple reaction time. Q. J. Exp. Psychol. 65, 151–164 (2012).

59. Ishai, A. & Sagi, D. Common mechanisms of visual imagery and perception. Science 268, 1772–1774 (1995).

60. Pearson, J., Rademaker, R. L. & Tong, F. Evaluating the mind’s eye: The metacognition of visual imagery. Psychol. Sci. 22, 1535–1542 (2011).

61. Sherwood, R. & Pearson, J. Closing the mind’s eye: Incoming luminance signals disrupt visual imagery. PLoS One 5, (2010).

62. Chang, S. & Pearson, J. The functional effects of prior motion imagery and motion perception. 105, 83–96 (2018).

63. Dijkstra, N., Hinne, M., Bosch, S. E. & van Gerven, M. A. J. Between-subject variability in the influence of mental imagery on conscious perception. 9, (2019).

64. Dijkstra, N., Mazor, M., Kok, P. & Fleming, S. Mistaking imagination for reality: Congruent mental imagery leads to more liberal perceptual detection. Cognition 212, (2021).

65. Neige, C. et al. Unravelling the Modulation of Intracortical Inhibition During Motor Imagery: An Adaptive Threshold-Hunting Study. Neuroscience 434, 102–110 (2020).

66. Jeannerod, M. Neural simulation of action: A unifying mechanism for motor cognition. in NeuroImage vol. 14 (Academic Press Inc., 2001).

67. Jeannerod, M. & Decety, J. Mental motor imagery: a window into the representational stages of action. Curr. Opin. Neurobiol. 5, 727–732 (1995).

68. Oldfield, R. C. The assessment and analysis of handedness: The Edinburgh inventory. Neuropsychologia 9, 97–113 (1971).

69. New, B., Pallier, C., Brysbaert, M. & Ferrand, L. Lexique 2: A new French lexical database. Behavior Research Methods, Instruments, and Computers vol. 36 516–524 (2004).

