## Supplementary material for "Imagining and reading actions: towards similar motor representations": e.g. Supplemental section

### Supplementary section

#### Material and method

##### Stimuli:

*Table 1: T-test results concerning the linguistic and psycholinguistic characteristics of stimuli*

| Factors | Flexion action verbs | Extension action verbs | T-tests |
|---|---|---|---|
| **Written frequency** | 10.24 | 8.58 | 0.77 |
| **Number of syllables** | 1.60 | 1.80 | 0.54 |
| **Number of characters** | 6.60 | 5.80 | 0.24 |
| **Spelling neighbors** | 4.40 | 4.60 | 0.92 |

*Table 2: Example of lexical stimuli used in the experiment.*

| **Flexion verbs** | **Extension verbs** | **Neutral stimuli** |
|---|---|---|
| je serre | j'étends | People |
| *I squeeze* | *I extend* | *People* |
| j'attrape | j'allonge | Jeudi |
| *I grasp* | *I stretch* | *Thursday* |
| j'agrippe | j'écarte | Appareil |
| *I grip* | *I spread* | *Device* |
| je presse | je tends | Écran |
| *I press* | *I hold out* | *Screen* |
| j'empoigne | je salue | Souris |
| *I clutch* | *I salute* | *Mouse* |

### Results

#### Percentage values

*Table 3: Percentage of correct responses for Control, Congruent, Incongruent and Trick conditions.*

| | Percentage of correct responses | | | |
|---|---|---|---|---|
| **Trials** | Control | Congruent | Incongruent | Trick |
| **Mean ±SD** | 52.69 ±11.06 | 60.16 ±10.52 | 53.68 ±14.37 | 51.90 ±10.94 |

#### Reaction times

To analyze all reaction times (correct and incorrect responses), we performed a repeated measures ANOVA with Congruence and Verb type (extension/flexion) as within-subject factors. No main effect was observed for Congruence ($F_{1,35}=0.580$, $p=0.451$, $\eta p^2=0.016$), Verb type ($F_{1,35}=0.029$, $p=0.864$, $\eta p^2=0.000$), nor interaction ($F_{1,35}=2.269$ $p=0.140$, $\eta p^2=0.060$). The same analysis on reaction times for correct answers yielded no main effect of Congruence ($F_{1,35}=2.553$, $p=0.119$, $\eta p^2=0.068$) nor Verb type ($F_{1,35}=0.845$, $p=0.364$, $\eta p^2=0.023$), and no interaction ($F_{1,35}=2.237$ $p=0.143$, $\eta p^2=0.060$) (See Figure 2 and raw data in Table 1).

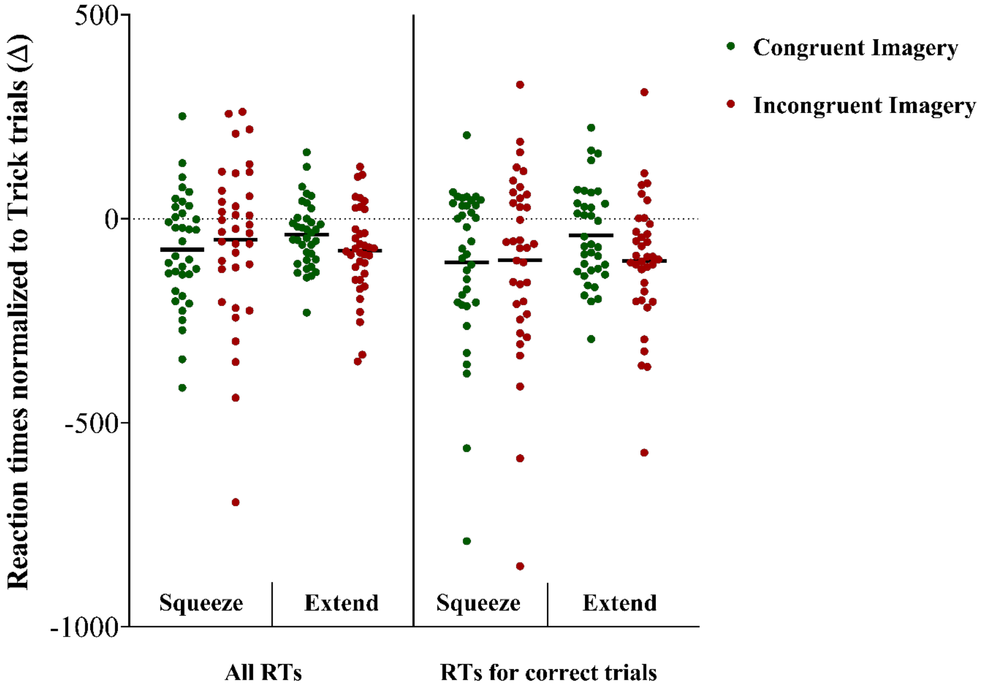


*Figure 3: Reaction times for Congruent and Incongruent normalized to Trick trials. The horizontal lines represent the mean of the group, while all circles represent individual values.*

*Table 4: Raw reaction times for all and correct responses in Congruent, Incongruent and Trick trials (in ms).*

| | All RTs | | | | | |
|---|---|---|---|---|---|---|
| | Squeeze verbs | | | Extend verbs | | |
| **Trials** | Congruent | Incongruent | Trick | Congruent | Incongruent | Trick |
| **Mean ±SD** | 1122.71 ± 311.70 | 1146.77 ± 337.70 | 1198.15 ± 364.74 | 1174.98 ± 318.58 | 1135.55 ± 301.40 | 1213.68 ± 334.15 |
| | **RTs for correct trials** | | | | | |

| Trials | Congruent | Incongruent | Trick | Congruent | Incongruent | Trick |
|---|---|---|---|---|---|---|
| **Mean ±SD** | 1106.32 ± 301.67 | 1112.30 ± 323.72 | 1213.43 ± 386.07 | 1190.21 ± 353.18 | 1127.42 ± 318.68 | 1231.09 ± 386.07 |

**EMGrms:**

The absence of muscular pre-activity during motor imagery was confirmed by a Friedman Anova revealing no significant difference in EMGrms before TMS artifact (100ms window prior the artifact) between rest (2.12±0.68μV), Congruent (2.49±0.95μV), Incongruent (2.38±0.89μV) and Control (Meaningless chain of consonants) (2.23±0.78μV) conditions ($p=0.31$; $r=0.01$).
